# Flexible reproductive seasonality in Africa-dwelling papionins is associated with low environmental productivity and high climatic unpredictability

**DOI:** 10.1101/2024.05.01.591991

**Authors:** Jules Dezeure, Julie Dagorrette, Lugdiwine Burtschell, Shahrina Chowdhury, Dieter Lukas, Larissa Swedell, Elise Huchard

## Abstract

At a time when seasonal cycles are increasingly disrupted, the ecology and evolution of reproductive seasonality in tropical vertebrates remains poorly understood. In order to predict how changes in seasonality might affect these animals, it is important to understand which aspects of their diverse patterns of reproductive phenology are linked to either the equally diverse patterns of rainfall seasonality (within-year variations) or instead the marked climatic unpredictability (year-to-year variations) occurring across the intertropical belt. Here, we gather birth and climatic seasonality data from 21 populations of 11 Africa-dwelling primate species from the papionin tribe, occupying a wide range of environments, including equatorial, tropical, temperate and arid climates. We investigate (1) the environmental variations that influence the intensity of reproductive seasonality, and (2) the reproductive stage that is synchronized with increased resource availability. Our results demonstrate wide variation in the intensity of birth seasonality between and within species. Across multiple measures of climatic variation, we found rainfall unpredictability to be the only clear predictor of the intensity of reproductive seasonality across populations, i.e., greater year-to-year variation in the amount of rainfall was associated with lower to no reproductive seasonality. Finally, we identified diverse patterns of reproductive phenology, with the most seasonal breeders generally aligning lactation with the peak in resource availability while other populations show more diverse patterns, where conception, lactation or weaning can all be synchronized with maximal food availability. This study sheds new light on the extent and ecological drivers of flexible reproductive phenology in long-lived tropical mammals, and may even contribute to our understanding of why humans give birth year-round.

## INTRODUCTION

Most animals face variation in their environment across the year (Boyce, 1979) in the form of seasonal fluctuations in rainfall, temperature and resource availability that affect their energy balance. Reproductive seasonality, the temporal clustering of reproductive events in the annual cycle, is thought to be beneficial because it synchronizes the most energetically costly reproductive stage with the seasonal food peak, thereby enhancing the condition and survival probability of mothers and offspring (Bronson, 2009; Bronson & Heideman, 1994). Variation in birth frequencies across the annual cycle is a continuous trait, ranging from a complete absence of reproductive seasonality (i.e., random distribution of births throughout the year), as in mountain gorillas (Campos et al., 2017), to cases in which all births occur within a few weeks each year, as in many lemurs (Wright, 1999).

Comparative studies investigating determinants of variation in reproductive seasonality across mammals have often been conducted at the level of the order (Rodents: Heldstab, 2021, Lagomorphs: Heldstab, 2021, ruminants: Rutberg, 1987; Zerbe et al., 2012, Carnivora: Heldstab et al., 2018, Primates: Di Bitetti & Janson, 2000; Heldstab et al., 2020; Janson & Verdolin, 2005) and thus focus on broad-scale macro-evolutionary patterns. These studies have typically detected a relationship between geographic latitude and birth seasonality, suggesting that at higher latitudes, birth seasonality is more pronounced, with a more intense birth peak (a birth peak being the temporal period in the annual cycle during which most birth occur). However, important gaps remain in our understanding of the determinants of reproductive seasonality. Few studies have attempted to quantify the extent of variation in reproductive seasonality across multiple populations of the same species (but see in African wild dogs (*Lycaon pictus*): McNutt, Groom, & Woodroffe, 2019, and in red-tailed monkeys (*Cercopithecus ascanius*): Struhsaker, 1997), or across closely related species sharing relatively similar diets, body sizes and life histories (but see in several ungulate species: Pereira, Dos Santos Zanetti, & Furlan Polegato, 2010; Spinage, 1973; Brogi et al., 2022; macaque species: Trébouet, Malaivijitnond, & Reichard, 2021). Thus, studies that control for major sources of variation in life history or broad dietary categories should be particularly useful for identifying the climatic drivers of variation in reproductive phenology.

The well-known association between latitude and reproductive seasonality fails to explain the diversity of reproductive seasonality patterns observed within restricted latitudinal ranges, such as in the tropics (Heldstab et al., 2020; Janson & Verdolin, 2005). In addition, latitude encapsulates multiple components of climatic variation, which need to be disentangled in order to identify the main climatic factors at play (Burtschell, Dezeure, Huchard, & Godelle, 2023). First, latitude correlates positively with the degree of environmental seasonality, measured as the magnitude of within-year variation (such as the difference between maximal and minimal monthly rainfall in the annual cycle) (Botero, Dor, McCain, & Safran, 2014). Further, latitude covaries negatively with environmental productivity, i.e., overall food availability in a given environment. Variation in productivity may alter the benefits of seasonal breeding, as populations living in more productive habitats may face less pressure to breed seasonally (Burtschell et al., 2023).

Finally, environmental predictability, independently of latitude and seasonality (Tonkin, Bogan, Bonada, Rios-Touma, & Lytle, 2017), could also influence breeding schedules. In locations with intense year-to-year environmental variation, a flexible reproductive phenology (i.e., individual ability to start a reproductive cycle at different timings of the year, in response to internal or external factors) may be more advantageous than a strictly seasonal reproduction (Brockman & van Schaik, 2005a; van Schaik & van Noordwijk, 1985). Indeed, regular delays or decreases in the food peak may lead to reproductive failures in strict seasonal breeders, thus reducing the fitness benefits of breeding seasonally. However, few studies have investigated the effects of environmental unpredictability on the intensity of reproductive seasonality, with mixed results so far. While English, Chauvenet, Safi, & Pettorelli (2012) found that higher inter-annual variation in food availability decreased the intensity of birth synchrony across 38 ungulate species, two studies of red deer, *Cervus elaphus* L. (Loe *et al*., 2005) and chacma baboons, *Papio ursinus* (Dezeure *et al*., 2023) found no effect of environmental unpredictability on reproductive seasonality. A recent modelling study similarly detected limited effects of environmental unpredictability on evolutionary transitions to nonseasonal breeding (Burtschell et al., 2023).

Aside from the selective pressures favouring a flexible reproductive phenology, relatively little is known about how birth timing varies in relation to the annual resource peak in long-lived species. In short-lived species, the full reproductive cycle, from conception to offspring nutritional independence (such as weaning in mammals or fledging in birds), can take place within a single productive season (Bronson, 2009). However, this is not the case for long-lived species, in which multiple stages of a female’s reproductive cycle can be aligned with the annual food peak, with varying fitness consequences (Dezeure et al., 2021). For example, females of some species may have to reach a certain threshold of body condition for the onset of reproduction and conception to take place (Brockman & van Schaik, 2005a), meaning that most conceptions are expected to follow a peak of food availability (Brockman & van Schaik, 2005a). In some other species, females may instead synchronize the costliest part of their reproductive cycle with the most productive season so as to enhance maternal condition and survival (Bronson, 2009; Bronson & Heideman, 1994), such that early-or mid-lactation occurs during the annual food peak, as in many primates (J. Altmann, 1980; Brockman & van Schaik, 2005a; Janson & Verdolin, 2005). Lastly, weaning is a critically vulnerable life stage, where juveniles must begin to forage for themselves (J. Altmann, 1980; Lee, 1996). Accordingly, several species have been shown to time their births so as to align weaning with the seasonal food peak (Brockman & van Schaik, 2005a; Janson & Verdolin, 2005), as occurs in most lemurs (Wright, 1999). Overall, the reasons underlying the observed variation in alignment of reproductive stages with the food peak across species and populations remain largely unknown.

In this study, we attempt to address the above gap in our understanding by investigating the evolutionary determinants of the intensity and timing of reproductive seasonality in Africa-dwelling papionin monkeys. We focus on papionins for several reasons. First, they exhibit relatively similar body sizes (large-bodied), life history traits (slow) and diet (mostly omnivorous) (Kingdon et al., 2012; Swedell, 2011), allowing us to investigate environmental effects on reproductive seasonality while controlling for these – potentially confounding – factors. Second, this taxonomic group displays a wide diversity of patterns of reproductive seasonality. Indeed, most baboon (*Papio* spp.) species are non-seasonal breeders (Bercovitch & Harding, 1993; Swedell, 2011) despite exhibiting variation in monthly birth frequencies (Cheney et al., 2004; Lycett, Weingrill, & Henzi, 1999), while mandrills (*Mandrillus sphinx*) (Setchell, Lee, Wickings, & Dixson, 2002), Kinda baboons (*Papio kindae*) (Petersdorf, Weyher, Kamilar, Dubuc, & Higham, 2019) and most mangabey species (i.e., *Cercocebus* and *Lophocebus* spp.) (Swedell, 2011) are seasonal breeders. Third, this species constellation exhibits great ecological flexibility, inhabiting arid areas, woodland savannahs, equatorial forests, and high altitude grasslands (J. Fischer et al., 2019; Kingdon et al., 2012; Swedell, 2011) (see also Figure 1). Fourth, within the African members of this tribe, baboons are one of the most well studied primate taxon, with data available from multiple populations of some species within the genus *Papio* (J. Fischer et al., 2019). Lastly, species from this taxonomic group, and in particular from the genus *Papio*, possess a variety of features shared with early hominins (Alberts et al., 2005; Brockman, 2005; Jolly, 2001): they are large, terrestrial and eclectic omnivorous primates (Alberts et al., 2005; Rhine, Norton, Wynn, & Wynn, 1989) that, unlike the great apes, have colonized African savannahs (Bobe, Martínez, & Carvalho, 2020) and give birth to a single offspring every one to three years (J. Altmann & Alberts, 2005; Swedell, 2011). Investigating the environmental determinants of their reproductive seasonality may thus shed new light on the evolution and maintenance of non-seasonal breeding in early hominins (King, 2022).

**Figure 1:**
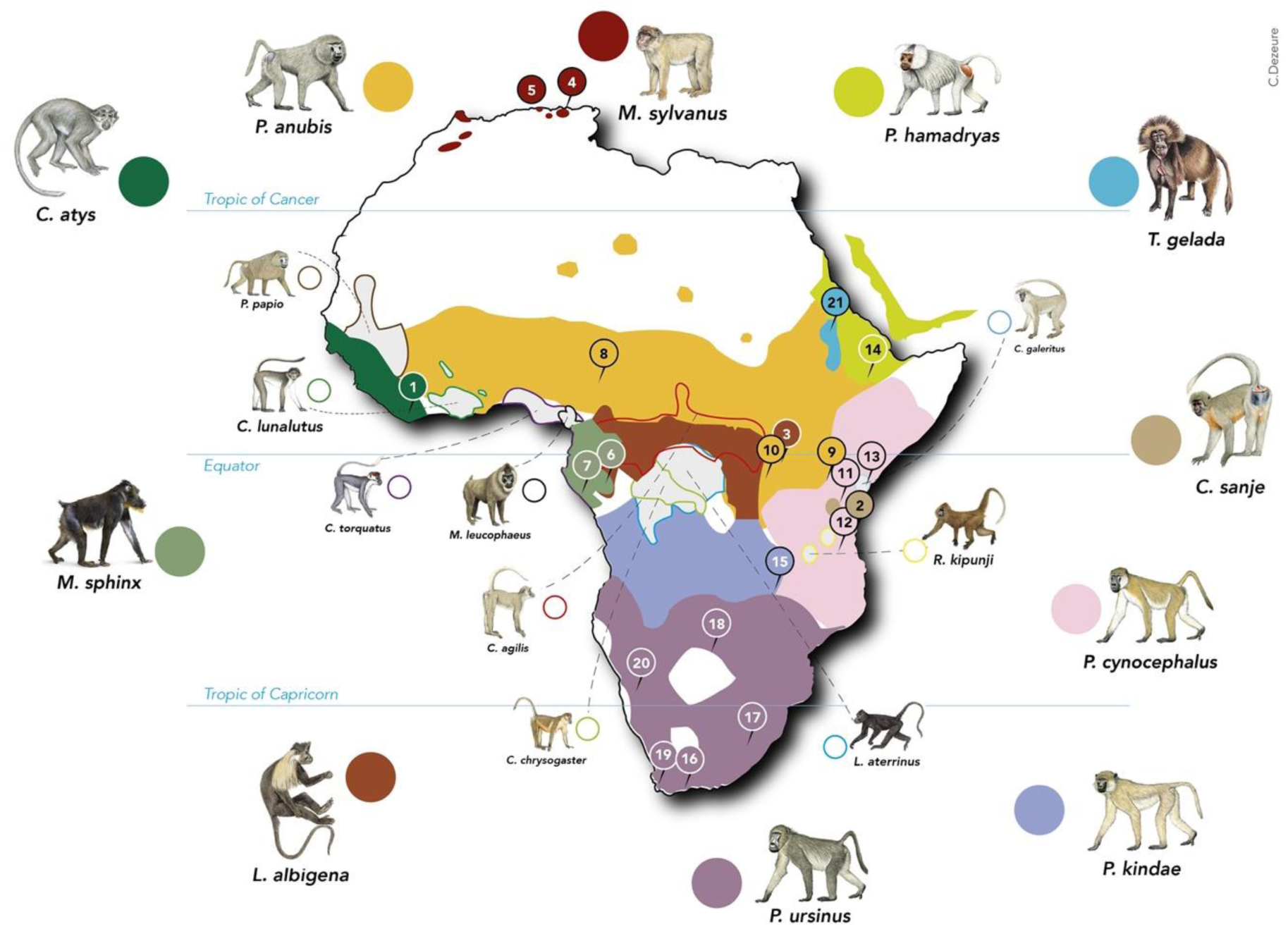
Distribution of the sampled species in Africa, and locations of the populations considered in this study. The species sampled in this study are depicted with relatively larger icons and species names, as well as full coloured circles. Species for which we could not find birth seasonality data are represented with smaller icons and names, as well as empty circles. The coloured areas on the map, corresponding to each species as indicated by the coloured circles, show the geographical distribution of each species. Within each range, small circled numbers show the location of the populations included in this study. 1: Taï, 2: Udzungwa Mountains, 3: Kibale, 4: Akfadou, 5: Tigounatine, 6: Lékédi. 7: Moukalaba-Doudou. 8: Gashaka-Gumti. 9: Gilgil. 10: Queen Elizabeth. 11: Amboseli. 12: Mikumi. 13: Tana River. 14: Filoha. 15: Kasanka. 16: De Hoop. 17: Drakensberg. 18: Moremi. 19: Tokai. 20: Tsaobis. 21: Simien. The species distribution ranges and icons come from Julia Fischer et al., 2017; Kingdon et al., 2012.

Here we ask three main questions regarding reproductive seasonality in the papionins in our sample:

i. What is the extent of inter- and intra-specific variation in patterns of reproductive seasonality, specifically regarding the height and width of the birth peak, as well as its timing in the annual cycle?
ii. What are the main environmental factors responsible for variation in the intensity of reproductive seasonality? We isolated eight components of environmental variation: latitude, environmental productivity, magnitude of seasonal variation in rainfall, number of rainy seasons, breadth of the rainy season, amount of between-year (unpredictable) variation in rainfall, between-year variation in the timing of the rainfall season, and the type of habitat. We tested the eight corresponding hypotheses (H1.1-1.8) and their associated predictions, which are listed in Table 1.
iii. In seasonally breeding populations, which stage of the reproductive cycle is synchronized with the food peak? We tested whether females match the seasonal food peak with conceptions (H2.1 -the ‘conception hypothesis’), lactation (H2.2 - the ‘lactation hypothesis’), or weaning (H2.3 - the ‘weaning hypothesis’).

**Table 1:**
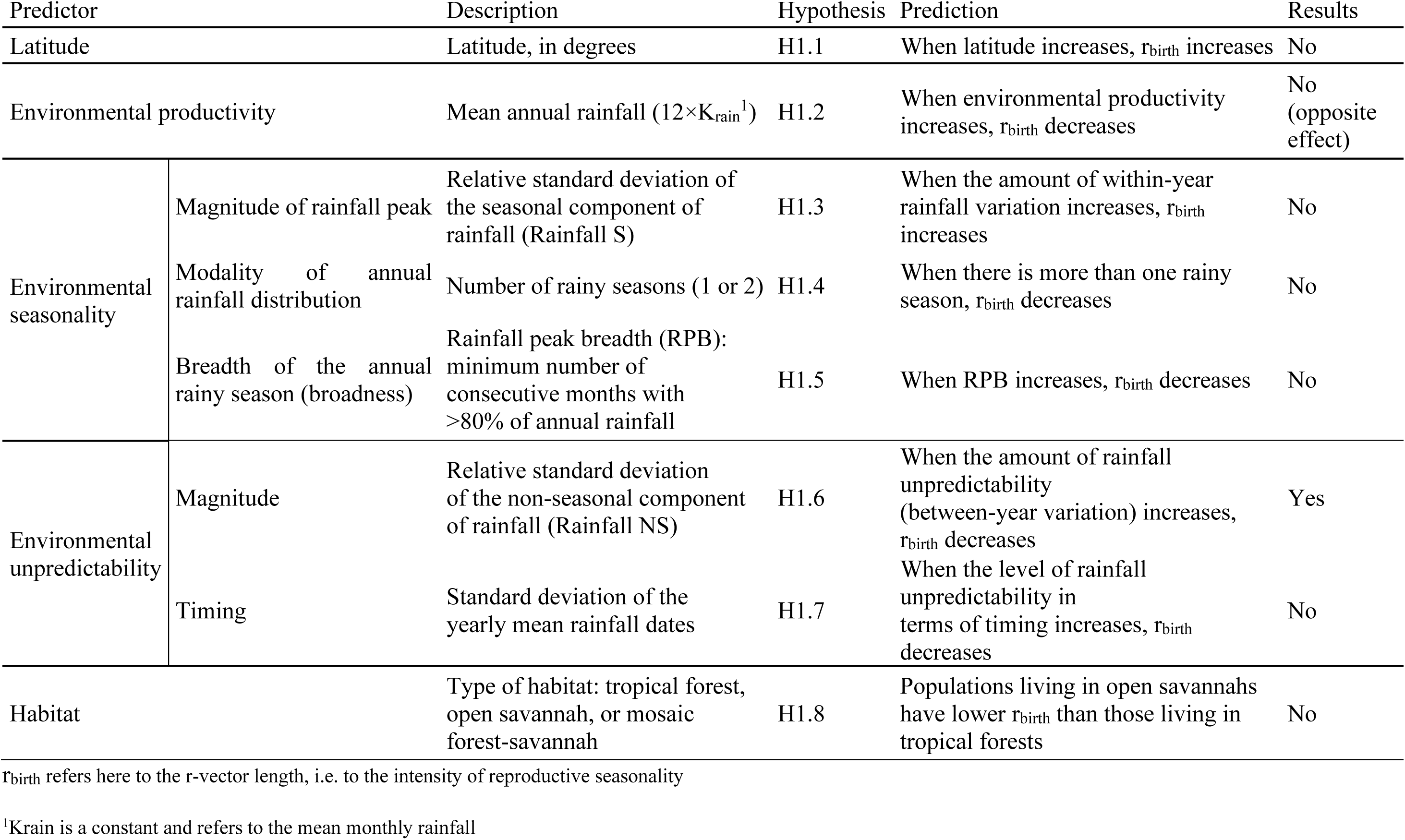
Hypotheses and predictions proposed on the effects of various environmental components on the intensity of reproductive seasonality

## METHODS

### 1- Sample and data selection

Our data set includes published reports on reproductive seasonality in natural populations of Africa-dwelling papionins from the genera *Cercocebus*, *Lophocebus*, *Macaca, Mandrillus*, *Papio*, *Rungwecebus* and *Theropithecus* (Figure 1). We selected papers that reported the number of births per month (except for yellow baboons, *Papio cynocephalus*, from Mikumi National Park, where births were provided in three month-periods). We obtained birth seasonality data from 21 wild populations representing 11 species: see Table S1 for references associated with each population, and Figure 1 for their locations. We did not find any monthly birth data for nine species of interest for which only data from captivity were available (Kingdon et al., 2012; Swedell, 2011): five species of *Cercocebus* (*agilis*, *chrysogaster*, *galeritus*, *lunulatus*, *torquatus*), as well as *Lophocebus aterrimus*, *Mandrillus leucophaeus*, *Papio papio* and *Rungwecebus kipunji* (Figure 1).

### 2- Birth seasonality data

We were interested in quantifying two components of reproductive seasonality in each population: (1) the mean population birth date, i.e. describing when most births mainly occur during the year, and (2) the intensity of population birth seasonality, i.e. describing how seasonal the births are. For our analysis, given the heterogeneity of the data (in some datasets, precise birth dates were available, but in most datasets, we could only obtain a count of births per month), we considered each birth to have occurred in the middle of the month (i.e., the 15^th^ of each month, except for February where births were considered to occur on the 14^th^). For the Mikumi population, we considered that births occurred in the middle of each 3 month-period. We then used a circular statistic and represented each birth event on the annual circle by a vector of length 1 and of angle θ representing its date (15×2×π/365.25 for January, (14+31) ×2×π/365.25 for February, etc.). We computed the mean vector (r-vector) per population, whose angle (converted in a date: µ_birth_) indicates the mean day of the year in which births occur (see Table S1) and is thus a measure of birth seasonality. We computed µ_birth_ using the function ‘circ.summary’ from the ‘CircStats’ package (Agostinelli & Lund, 2018). For populations with a significant birth peak, µ_birth_ represents the date of the population birth peak, i.e. when births are the most likely to occur in the annual cycle. The length (r_birth_) of the r-vector measures the intensity of birth seasonality, i.e., the degree of uniformity of the birth distribution across the annual cycle, varying from 0 to 1 (Di Bitetti & Janson, 2000; Janson & Verdolin, 2005; Thompson & McCabe, 2013). When r_birth_=0, births are evenly spread across months (i.e., non-seasonal), while when r_birth_=1, births all occur during the same month of the year (extremely seasonal). After comparing several classical measures of reproductive seasonality (Supplementary Materials, Appendix S1), we used only r_birth_ to measure the intensity of reproductive seasonality, as this measure is more robust to differences in sample size than other metrics, facilitating the comparison of seasonality measures between populations (Janson & Verdolin, 2005; Thel, Chamaillé-Jammes, & Bonenfant, 2022).

### 3- Environmental data

#### i. Two indicators of environmental variation: rainfall and NDVI

In order to test our set of hypotheses, we considered environmental variation through two components: rainfall and the Normalized Difference Vegetation Index (NDVI). NDVI produces a quantitative index of vegetation productivity, where higher values indicate a higher degree of vegetation cover (Didan, Barreto Munoz, Solano, & Huete, 2015). Climatic seasonality in Africa (and in most tropical habitats) is mainly characterized by within-year variation in rainfall (Alberts et al., 2005; Feng, Porporato, & Rodriguez-Iturbe, 2013; Van Schaik, Terborgh, & Wright, 1993), which has been successfully used as an indicator of food availability for several of our studied populations (Alberts et al., 2005; Hill, Lycett, & Dunbar, 2000; Petersdorf et al., 2019; Tinsley Johnson, Snyder-Mackler, Lu, Bergman, & Beehner, 2018). Yet, NDVI values have to be used with caution when comparing productivity across environments (Pettorelli et al., 2005), which is why we opted to use rainfall to test hypotheses H1.1-H1-8. For example, the mean annual NDVI value at De Hoop, one of our driest habitats, was almost equal to that of Gashaka, one of our wettest habitats. We thus used variation in rainfall, rather than in NDVI, to disentangle the various components of climatic variation that may affect the intensity of reproductive seasonality (such as environmental productivity, predictability and seasonality) when testing hypotheses H1.1-H1.8.

However, we opted to use NDVI as an index of food availability to calculate the timing of the food peak when testing hypotheses H2.1-H2-3, as Africa-dwelling papionins rely mainly on plants for their diet (Swedell, 2011). Specifically, the annual peak in NDVI – and presumably in food availability - is likely to lag behind that of rainfall by a few weeks, and this lag duration may vary depending on local climatic or environmental conditions (Bercovitch & Harding, 1993; Dezeure et al., 2021; Jarvey, Low, Pappano, Bergman, & Beehner, 2018). Consequently, we thought that using NDVI would be preferable than rainfall to assess the timing of the annual food peak, and test hypotheses regarding its match to the birth peak in each population. Nevertheless, the use of NDVI as an index of food productivity in papinions, which are not herbivorous (except for Theropithecus gelada), could be arguable, and results of this analysis would be discussed accordingly.

#### ii. Data extraction

Daily rainfall was extracted from satellite data sensors with the Giovanni NASA website (product TRMM 3B42) (Huffman, Bolvin, Nelkin, & Adler, 2016) using a 0.25×0.25 degree resolution (corresponding to between 28×28km at the equator and 23×23km at 35° latitude). The GPS coordinates used for this extraction are indicated per population in Table S2, and were assessed either from indications about the home ranges of the habituated groups per population when available in the literature, or alternatively from the geographical location (Park, Reserve or nearby city) of the population (see also Figure 1). Monthly cumulative rainfall (summed across daily values) was subsequently computed between January 1998 and December 2019. We therefore gathered 22 years of rainfall data per population over the same period of time.

We then extracted the mean NDVI per 16 day-period on a 500m × 500m resolution within the same geographical areas used for rainfall extraction (see GPS coordinates in Table S2) between March 2000 and March 2017 (data before and after these dates were not available at the time of data extraction) using MODIS data (MODIS13A1 product) provided by NASA (Didan et al., 2015). Daily NDVI was computed by linear interpolation and then averaged to obtain a monthly value across 18 years.

#### iii. Components of environmental variation

In order to test our hypotheses, we identified multiple components of rainfall variation within and across years. First, we decomposed for a given site the observed rainfall value into three components as follows: Rainfall_m,i_ = K_rain_ + Rainfall S_m_ + Rainfall NS_m,i_, where m is the month of the year (going from January to December) and i is the year (from 1998 to 2019). K_rain_ is a constant, equalling the mean monthly rainfall across 22 years of records (Figure S1). Rainfall S_m_ is the seasonal component of rainfall, i.e., the rainfall value, averaged across 22 years, for each month of the year, minus K_rain_ (Figure S1). For example, for a given site, Rainfall S_1_ (m=1) is the mean of all January rainfall values. The term Rainfall S thus captures the seasonal component of rainfall variation in the annual cycle, i.e., its within-year variation. Finally, Rainfall NS _m,i_ is the non-seasonal component of rainfall, i.e., the difference between the observed rainfall value in any month at a given site (Rainfall_m,i_) and the predictable component of rainfall variation for that particular site in that particular month (K_rain_ + Rainfall S_m_) (Figure S1). This captures the unpredictable, i.e., between-year, rainfall variation. Using these measures, we assessed the following for each population (see Table S2 for the values associated with each population):

- Environmental productivity, or mean annual rainfall, equal to 12×K_rain_.

- Magnitude of environmental seasonality. We computed the magnitude of within-year rainfall variation, as the relative standard deviation (SD) of Rainfall S standardized for environmental productivity, given by the formula: 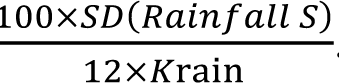. The higher the rain value, the more seasonal is rainfall variation.

- The number of rainy seasons per year. Using predictable rainfall variation (K_rain_ + Rainfall S), we assessed graphically, for each population, the number of rainy seasons per year.

- The length of the rainy season. For environments with only one rainy season, we further calculated the rainfall peak breadth (RPB), which is the minimum number of consecutive months of the year during which 80% of the annual rainfall (12*K_rain_) occurs. This measure is meaningless for environments with more than one rainy season, and we thus excluded the populations living in such environments from this analysis.

- Magnitude of environmental unpredictability. We computed the magnitude of between-year rainfall variation, as the relative standard deviation of Rainfall NS, standardized by environmental productivity, given by the formula: 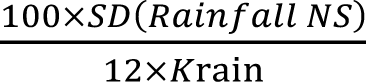. The higher the rain value, the more unpredictable rainfall variation is.

Using the literature, we categorized the habitat of each population into three types: tropical forest, open savannah, and mosaic forest-grassland (Table S2).

In addition to the magnitude of environmental unpredictability, we were interested in quantifying unpredictability in the timing of the annual rainfall peak, i.e., quantifying how much the timing of rainfall varied between years. Details on the procedure can be found in Appendix S2, but briefly, using circular statistics, we computed standard deviations of the mean dates of the annual rainfall peak over the 20 years sampled. Values close to zero mean that rainfall peak occurs the same month every year, and the higher the value, the more variation in rainfall peak’s timing between years.

Lastly, we computed the mean monthly NDVI across 18 years (i.e., K_NDVI_) and the seasonal component of NDVI variation for each month of the year (i.e., NDVI S), following the same notation used to disentangle the components of rainfall variation. To characterise the timing of the seasonal food peak, we used circular statistics to compute the mean annual NDVI date, µ_NDVI_ (see Appendix S2 for methodology and Table S2 for values of µ_NDVI_).

#### iv – Phylogenetic tree

We used the branch length of Version 3 of the 10kTreesPrimates consensus tree (Arnold, Matthews, & Nunn, 2010). Two species of interest were absent from this tree: *Cercocebus sanjei* and *Papio kindae*. For the former, we substituted *Cercocebus galeritus*, its closest relative. For the latter, following recent genetic studies (Jordan et al., 2018; Rogers et al., 2019), we added *Papio kindae* in the same branch as *Papio ursinus*, using the function ‘bind.tip’ from the package ‘phytools’ (Revell, 2012).

### 4- Statistical analysis

All statistical analyses were conducted in R version 3.5.0 (R Core Team, 2019).

#### i. Factors affecting the intensity of reproductive seasonality

We considered eight potential environmental parameters associated with reproductive seasonality, which are listed along each corresponding prediction in Table 1.

For each hypothesis, we plotted r_birth_ (the r-vector length measuring the intensity of reproductive seasonality in a given population) versus the tested predictor. We then checked the significance of the relationship between each predictor and r_birth_ while controlling for phylogeny using Bayesian phylogenetic generalized linear mixed models, with a Beta regression and a logit link function, with the package ‘brms’ (Bürkner, 2017). We included the phylogenetic relationship between species as a covariance matrix, which was derived from the phylogenetic tree.

Given the low number of populations in this study, and the collinearity between some of our environmental parameters (multivariate models had variance inflated factors >3), we were not able to run stable multivariate models. For example, the magnitude of environmental seasonality was negatively associated with the duration of the rainy season (cor=-0.94, t=-9.15, p<10^-4^), and environmental productivity was negatively associated with both latitude (cor=-0.55, t=-2.69, p=0.016) and the magnitude of environmental unpredictability (cor=-0.80, t=-5.48, p<10^-4^).

Therefore, our Beta regressions included as a response variable r_birth_ values, one fixed effect (each environmental predictor in turn listed in Table 1, standardised if continuous) and the phylogenetic matrix as a random effect. Given the high variation in sample size (i.e., number of births recorded) between populations, we used a weighed regression, where the weight given to each data point equals to log(N) / minimum(log(N)) so that the population with the lowest sample size counts for 1 observation, and the other populations count for more observations depending on their sample size, following a logarithmic scale ; the logarithmic scale was chosen to account for the diminishing return of increasing the sample size of samples that are already large. Beyond a given sample size, further increases in sample size do not affect much r_birth_ estimates, which are already stable and precise. For each model, we set an informative prior and used 3 000 iterations, a burn-in of 1 000 and 3 chains. We visually inspected for convergence and checked the absence of autocorrelations for the posterior distributions of fixed and random effects. The predictors were considered statistically significant when their associated 95% confidence intervals did not cross 0.

Finally, we extracted the phylogenetic signal in our dataset with the metric of Blomberg’s K, allowing us to compare it with other signals from other traits. To do so, we computed the mean r_birth_ per species, and use the ‘phylosig’ function with 1000 simulations from the ‘phytools’ package (Revell, 2024).

#### ii. Timings of conceptions, births and weaning in relation with NDVI seasonality

We tested H2 only for those populations for which a significant birth peak can be detected, as it does not make sense to test which period of the reproductive cycle is matched with the annual food peak if there is not a clear seasonal pattern of births in one population. We therefore assessed whether each population had a significant birth peak using the Rayleigh test for circular statistics, more precisely the ‘r.test’ function from ‘CircStats’ package (Agostinelli & Lund, 2018). For each population, when the P-value associated with the Rayleigh test was <0.05, meaning that the null hypothesis of a uniform birth distribution could be rejected, the birth peak was considered significant. With this approach, some populations with relatively low reproductive seasonality (low r_birth_) but with a large number of births are included, which should be taken into account when interpreting the results. Among these populations with a significant birth peak (see Table S1), we investigated which reproductive stage (H2.1, ‘conception’; H2.2, ‘lactation’; or H2.3, ‘weaning’) was synchronized with the annual NDVI peak, i.e. µ_NDVI_. Additional details are given in Supplementary Materials, Appendix 3 & Table S3. We employed exact two-sample Fisher-Pitman permutation tests, using the ‘oneway_test’ function from the ‘coin’ package (Hothorn, Hornik, Van De Wiel, & Zeileis, 2006). This function tests if the observed monthly value of NDVI during a target period, which depends on the hypothesis tested, is significantly higher than monthly values of NDVI randomized across the entire year. For example, using mean gestation length to infer the annual distribution of conception dates, we tested H2.1 by asking if females tended to conceive during, soon before, or soon after the annual food peak, looking at seasonal NDVI values respectively in (i) the six months surrounding µ_conc_, (ii) the three months before µ_conc_ and (iii) the three months after µ_conc_.

## RESULTS

### 1) How variable are patterns of reproductive seasonality?

The annual distribution of births for each population is shown in Figure 2, alongside seasonal variation in rainfall and NDVI. The intensity of reproductive seasonality varies widely across species: *Papio hamadryas* (Filoha population: r_birth_ =0.02) and most *Papio anubis* populations (r_birth_ <0.22) show non-seasonal births while *Mandrillus sphinx* (Lékédi: r_birth_ =0.67 and Moukalaba-Doudou: r_birth_ =0.80 resp.) and *Papio kindae* (Kasanka: r_birth_ =0.50) exhibit pronounced birth seasonality (Figure 3). The phylogenetic signal associated with the intensity of reproductive seasonality is substantial (Blomberg’s K=1.83, pvalue<0.01), indicating that more closely related species have more similar patterns of reproductive seasonality. Among papionins, the genera *Mandrillus*, *Cercocebus* and *Macaca* show strong reproductive seasonality, whereas the genera *Lophocebus*, *Papio* and *Theropithecus* show an overall lower intensity of birth seasonality, which may be associated with greater flexibility within species (Figure 3). Such flexibility is particularly pronounced in *Papio ursinus* populations, extending from low (de Hoop: r_birth_ =0.10; Tokai: r_birth_ =0.22, and Tsaobis: r_birth_ =0.10) to moderate birth seasonality (Drakensberg: r_birth_ =0.42; Moremi: r_birth_ =0.37) (Figure 3), while intra-specific variation seems less marked for the other species represented by multiple populations in our sample.

**Figure 2:**
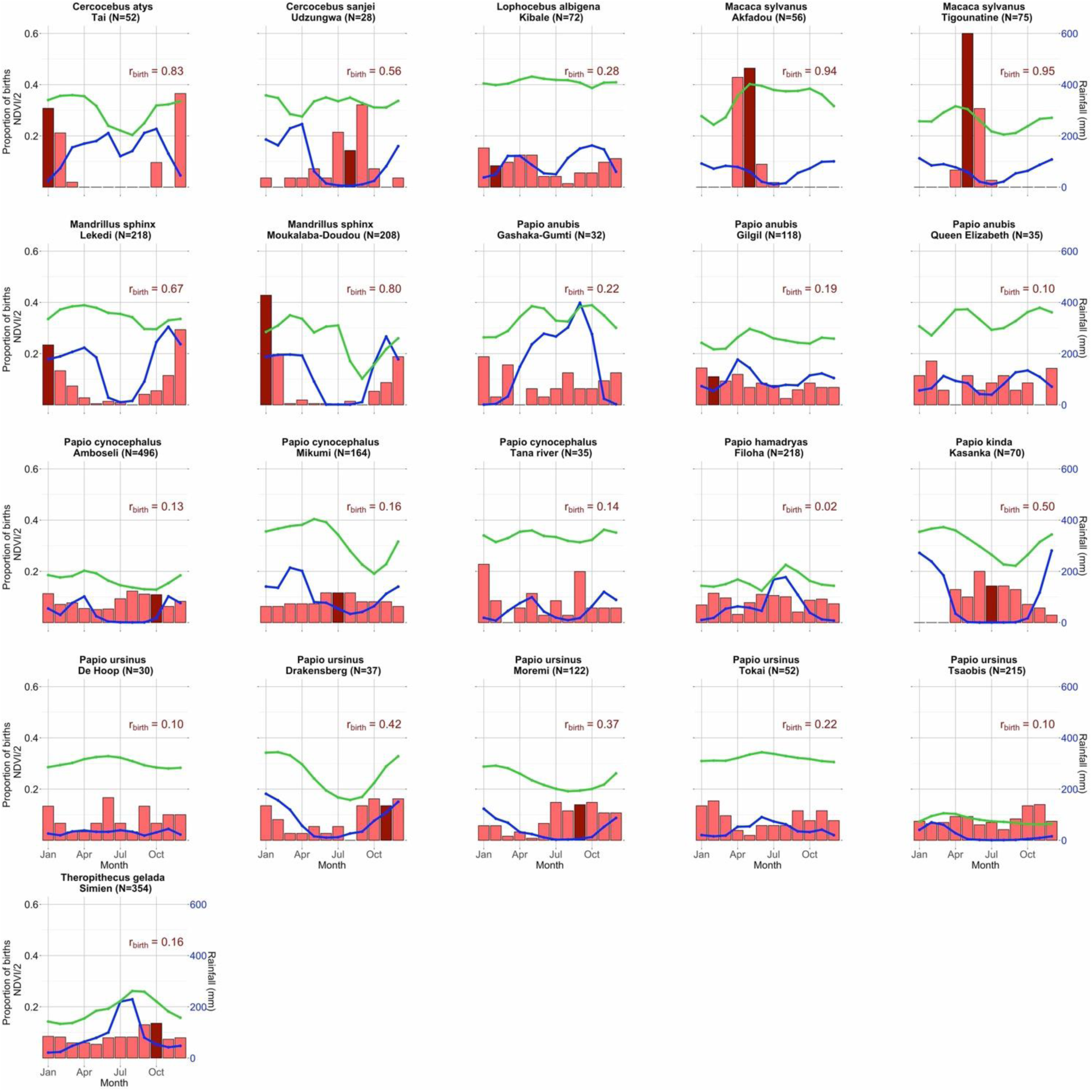
Monthly distribution of births in relation to rainfall and NDVI seasonality The proportion of births per month (left side of the y-axis) is represented with red bars. In addition, the darker red bar indicates the month of the mean birth date (µbirth) for seasonal breeding populations. We indicate in darker red the value of the r-vector length (r_birth_) for each population (top-right corner of each panel). We represent the mean monthly rainfall (equalled to Krain + Rainfall S, in mm) in blue (right side of the y-axis), and the mean monthly NDVI (equal to KNDVI + NDVI S and divided by 2 for graphical purposes) in green (left-side of the y-axis). The species and population names are indicated on top of each panel, along with the number of births observed (N).

**Figure 3:**
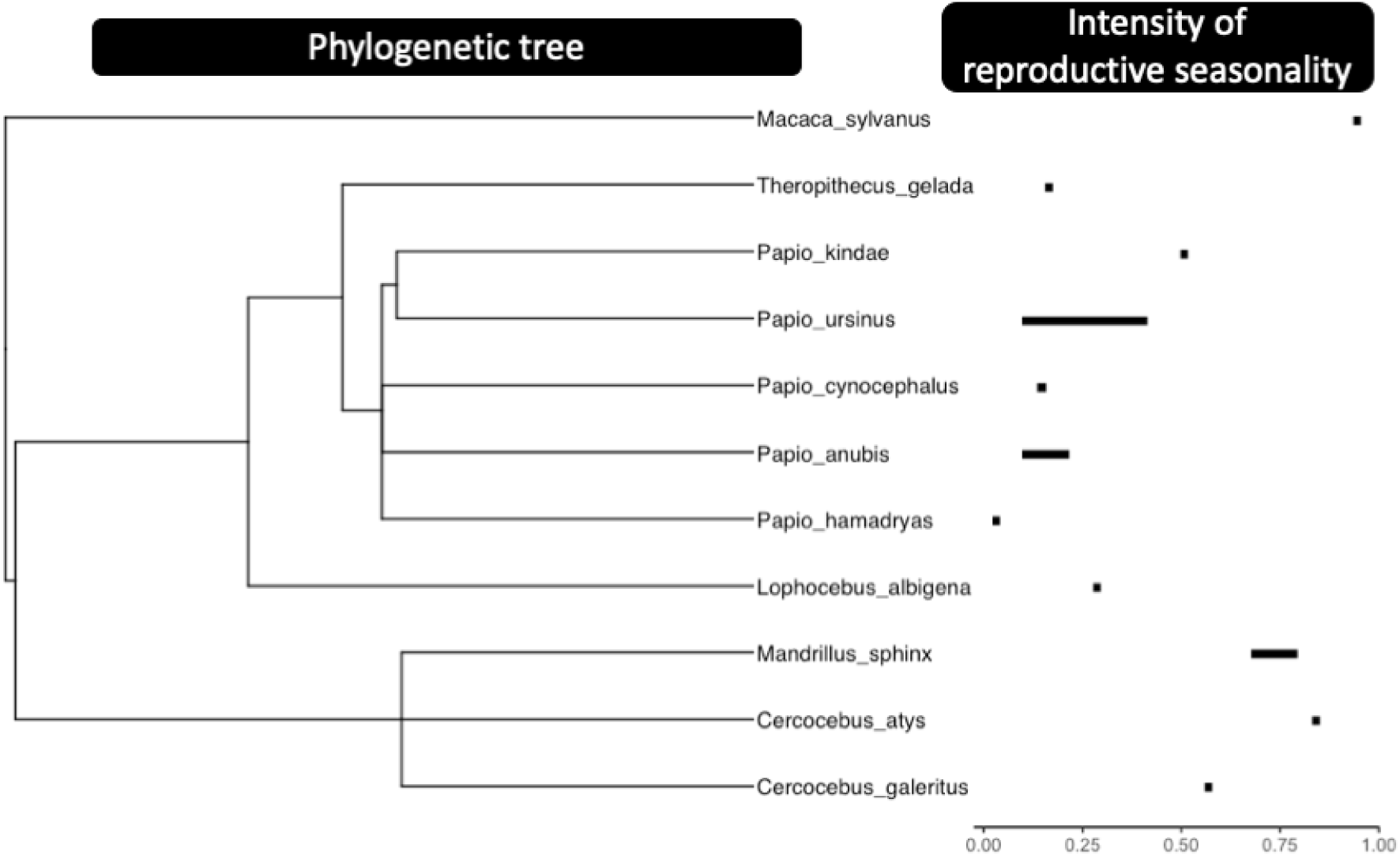
Phylogenetic tree of the studied species including the variation in the intensity of reproductive seasonality for each species. The intensity of birth seasonality is here quantified using the r_birth_ value. For those species represented by more than one population, the length of the black segment displays the variation in the intensity of birth seasonality among populations.

The timing of birth seasonality can also be surprisingly variable, even between species that live in adjacent geographical ranges: for instance, *Papio kindae* from Kasanka give birth mainly around July, whereas *Papio ursinus* from Moremi give birth primarily between July and November (Figure 2).

### 2) What are the ecological parameters correlated with the intensity of reproductive seasonality?

We detected a significant correlation between the magnitude of environmental unpredictability and the intensity of reproductive seasonality, while controlling for species relatedness, which supported our prediction H1.6 (Table 2): the higher the magnitude of between-year variation in rainfall, the lower the intensity of reproductive seasonality (Figure 4F). We also found an effect of habitat productivity that contradicted our prediction H1.2 (Table 2): the lower the mean annual rainfall, the lower the intensity of reproductive seasonality (Figure 4B). Lastly, we did not find any support for the other six hypotheses: there was no effect of latitude, magnitude of rainfall seasonality, number of rainy seasons, breadth of the rainfall peak, unpredictability in the timing of the annual peak of rainfall, or habitat on the intensity of reproductive seasonality (Table 2, Figure 4).

**Table 2:**
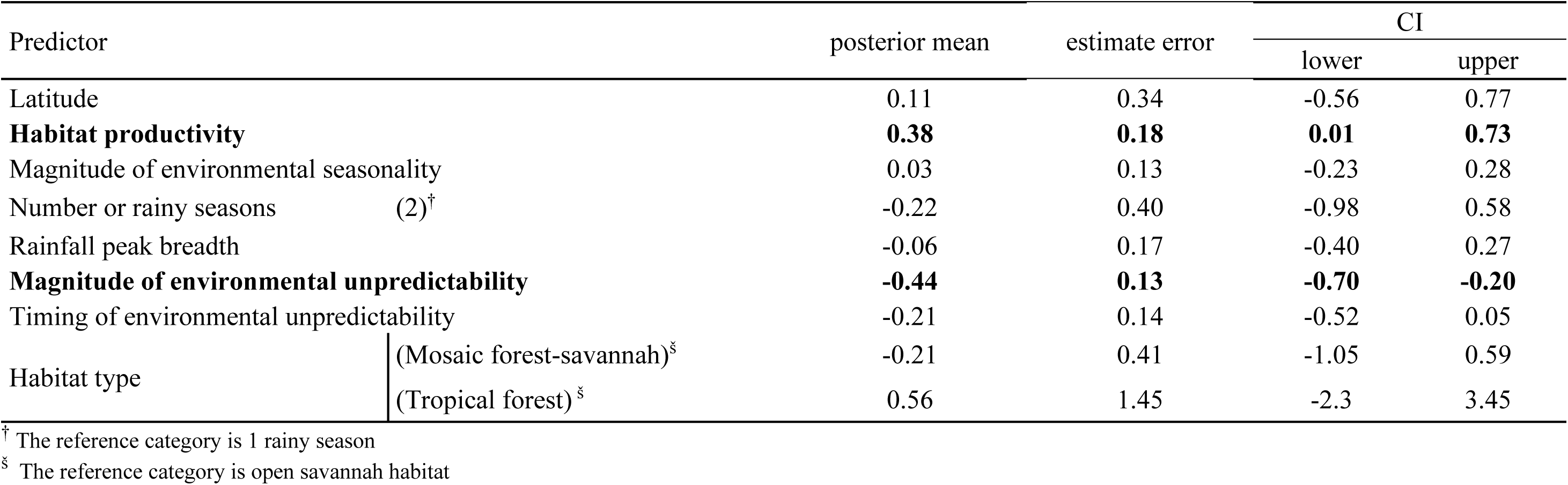
Influence of several components of rainfall variation on the reproductive seasonality of Africa-dwelling papionin populations The table shows the posterior mean, the estimate error and the 95% (marginal) confidence intervals (CI) associated for each posterior distribution of the predictors of the Beta regression brms models including species’ relatedness as random effect, r_birth_ as response variable, weighted by the log-transformed number of observations, and each predictor as the only fixed effect of a univariate model. Significant effects are indicated in bold. For categorical predictor, the tested category is indicated between parentheses.

**Figure 4:**
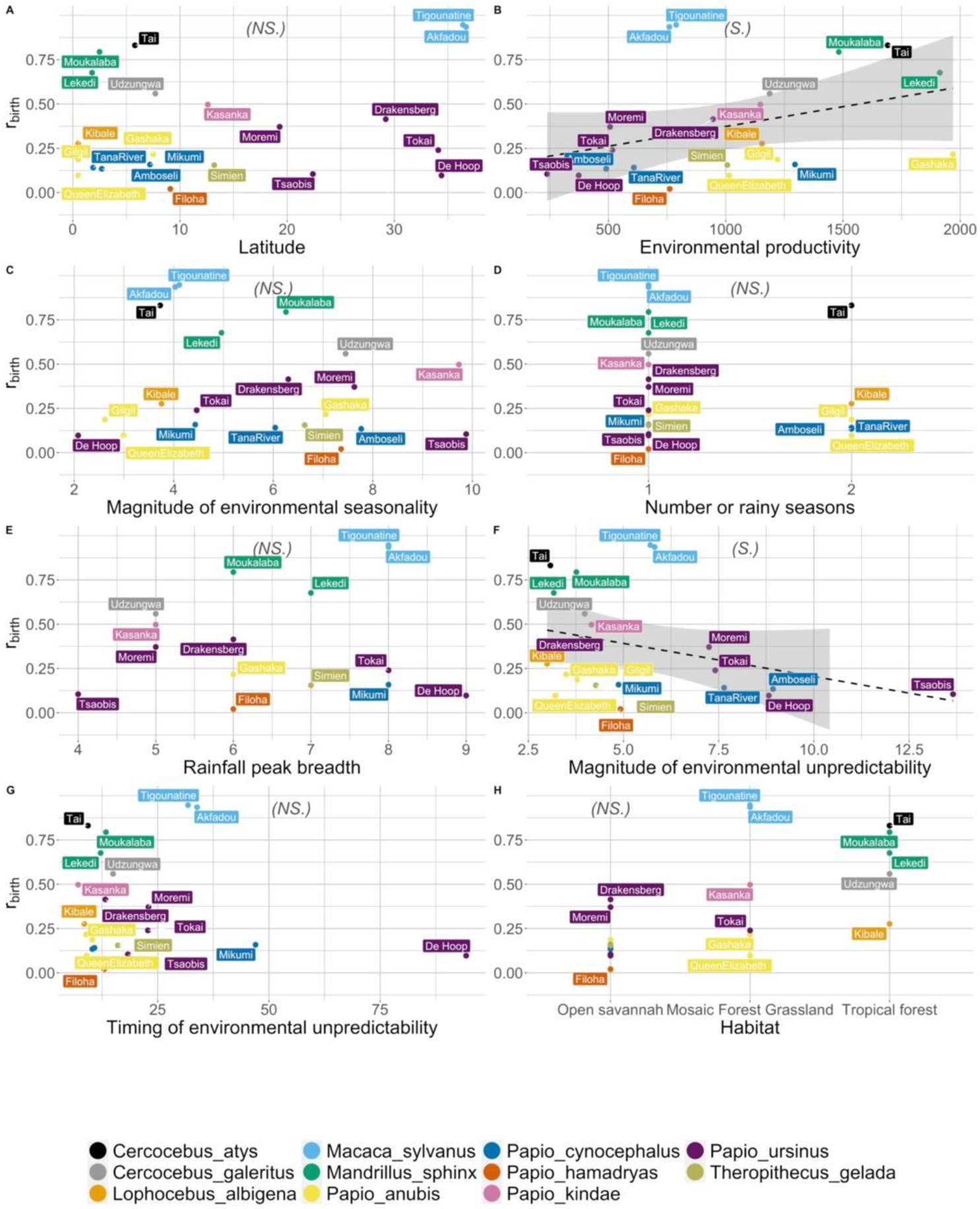
Effect of multiple environmental factors on the intensity of reproductive seasonality. We plotted the intensity of reproductive seasonality (r_birth_) depending on latitude (Panel A), environmental productivity (indexed by mean annual rainfall in mm, Panel B), the magnitude of environmental seasonality (i.e. of the rainfall peak, Panel C), the number of rainy seasons (Panel D), the rainfall peak breadth (RPB, Panel E), the magnitude of environmental unpredictability (i.e. of variation in the non-seasonal component of rainfall, Panel F), the timing of environmental unpredictability (i.e. between-year variation in rainfall timings, Panel G), and habitat type (Panel H). For each panel, each dot represents a population (with the population name annotated), and the colour indicates the species (see legend at the bottom). The dashed black line represents the linear regression, and the shaded area displays 95% confidence intervals. On top of each panel, we indicated in italic and between parentheses the significance of each predictor (NS for non-significant, S for significant).

### 3) Which stage of the reproductive cycle is timed with the annual food peak?

When considering all taxa in our sample as a whole, none of our three hypotheses were clearly supported (H2.1, H2.2, H2.3). However, in five of the six populations with the strongest reproductive seasonality (r_birth_ >0.5), females appeared to synchronize lactation with the annual NDVI peak, and overall lactation was generally aligned with the NDVI peak in 8 of the 14 populations with a significant birth peak (P-value of the Rayleigh test <0.05) (Table 3, Table S4). In less seasonally breeding populations (r_birth_ <0.5), females were more variable in the reproductive stages that were timed with the annual NDVI peak, ranging from conception (6 of 14 populations, and mainly before than after conception: e.g. Amboseli or Kibale) to weaning (3 of 8 populations: e.g., Moremi or Gilgil) to none (e.g., Simien) or all of these stages (e.g., Udzungwa) (Table 3, Table S4). Moreover, the timing of the annual NDVI peak compared to the mean conception, birth or weaning dates was highly variable between populations (Table S4).

**Table 3:**
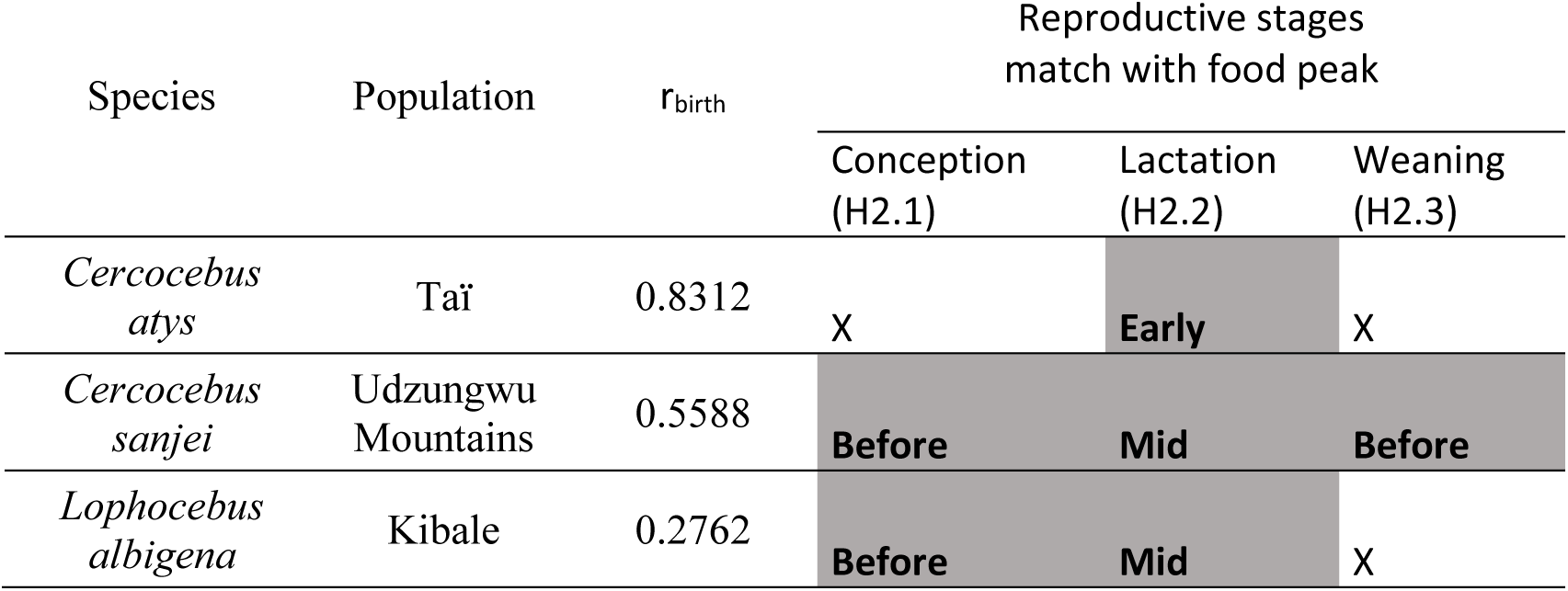
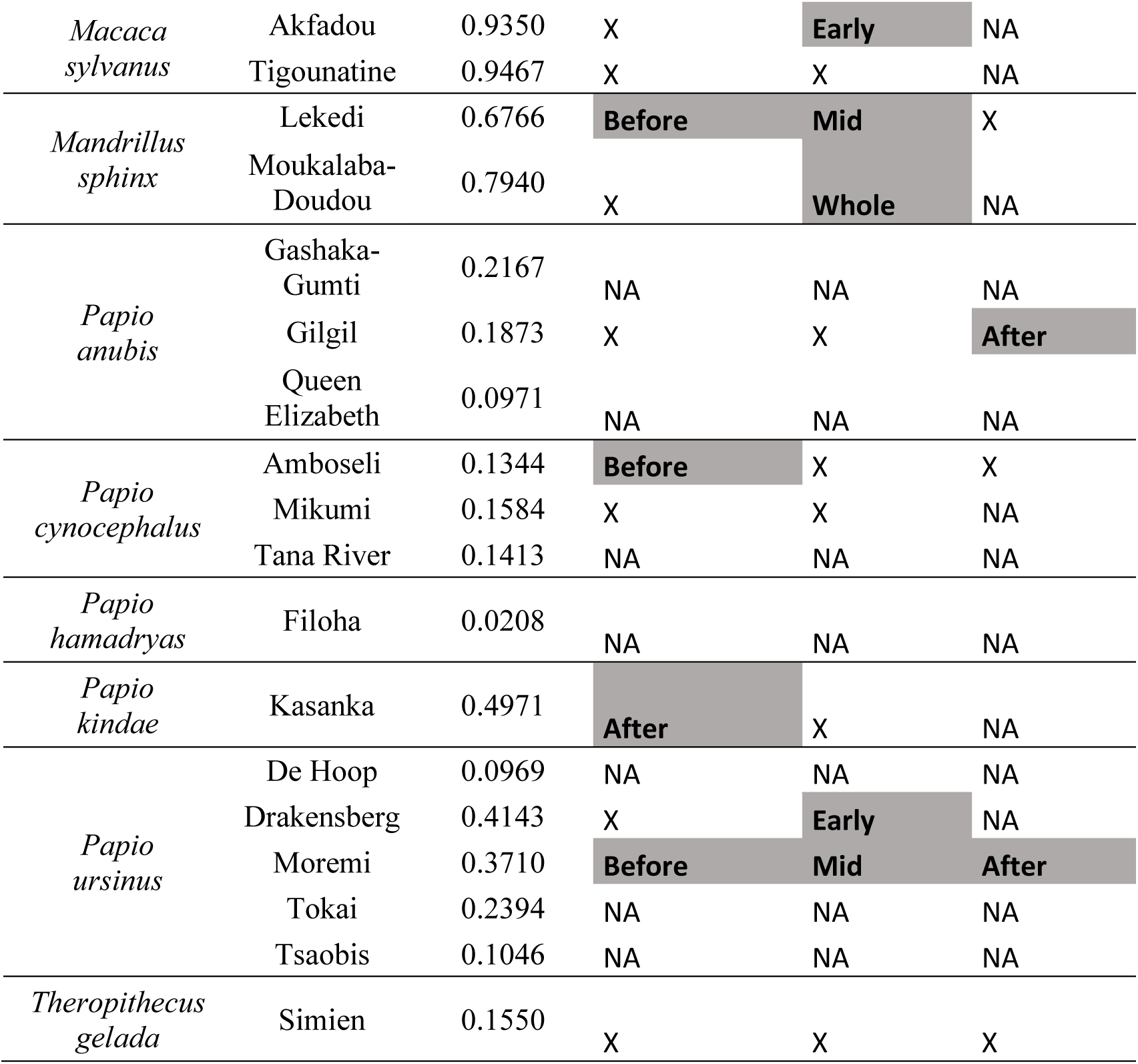
Reproductive stage matched with the food peak in sampled populations. Cells are filled with NAs when the alignment of the food peak with this reproductive stage was not tested (either because the data on weaning age in this population was missing, or because the birth peak of the population is non-significant, i.e., with P-value of the Rayleigh test >0.05). Cells are filled with X when there is no alignment, and shaded when there is an alignment of the given reproductive stage with the food peak.

## DISCUSSION

We revealed strong inter- and intra-specific variation in the intensity of reproductive seasonality as well as in the annual timing of births in Africa-dwelling papionins. Our study further emphasizes the importance of environmental unpredictability for the evolution of flexible reproductive seasonality. Lastly, we found that females from different populations of Africa-dwelling papionins match different reproductive stages with the annual food peak.

### Papionins exhibit flexible reproductive seasonality

Of the sampled *Papio* populations, most showed very little seasonality and *Papio kindae* departed from the overall baboon pattern in being relatively highly seasonal breeders. Such diverse patterns of reproductive seasonality within a single genus have rarely been reported outside primates (see for example *Cervus*: English et al., 2012; Loe et al., 2005; Rutberg, 1987, *Damaliscus*: Rutberg, 1987, *Ovis*: Rutberg, 1987, *Ursus*: Spady, Lindburg, & Durrant, 2007, *Mustela*: Heldstab et al., 2018, *Vulpes*: Heldstab et al., 2018), but apparently occur in some other primate genera with large distribution ranges, such as *Alouatta* (Di Bitetti & Janson, 2000; Janson & Verdolin, 2005), *Cercopithecus* (Heldstab et al., 2020; Janson & Verdolin, 2005), *Cebus* (Janson & Verdolin, 2005) and *Macaca* (Heldstab et al., 2020; Janson & Verdolin, 2005; Trébouet et al., 2021). Given the limited taxonomic scale of our study, it is impossible to establish whether seasonal breeding was the ancestral state in papionins, but it seems possible that the loss of seasonal reproduction is a derived state affecting the *Papio* genus (Fig 3). The estimated phylogenetic signal is significant and shows that among the 11 sampled papionin species, the intensity of birth seasonality is more similar among two closely related species. The value of this phylogenetic signal (Blomberg’s K=1.83) is relatively high, among the highest of many life history traits (such as age at maturity, adult mortality, clutch size in birds, sexual dimorphism, etc.), and higher than behavioural traits (such as daily movement distance, prey size, preferred body temperature, etc.), that are more labile (Blomberg, Garland, & Ives, 2003). Despite this strong impact of phylogeny on the intensity of reproductive seasonality, our study emphasizes the importance of the variations in birth seasonality between two closely-related species, or even within a single species.

Importantly, the key adaptive trait that evolved in the *Papio* genus may not be simply the loss of breeding seasonality *per se*, but the evolution of a flexible reproductive phenology. A same papionin female can give birth at different timings for successive birth events, depending on her own individual traits or physiological constraints, or alternatively depending on the strategies of other females in the same social group, as shown in *Papio ursinus* (Dezeure et al., 2021, 2023). This reproductive flexibility at the individual level necessarily shapes population patterns of reproductive seasonality, leading to lower reproductive seasonality. Reproductive flexibility at the population level could be defined as the ability for different populations of a same species to exhibit diverse patterns of reproductive seasonality, depending on the environmental conditions. Such flexibility is observed at the population level in Papio ursinus, living in a large distributional range characterized by exceptional ecological diversity, which includes cold and temperate climates, oceanic and mountainous ecosystems, and tropical and arid savannahs. Indeed, populations of this species exhibit a wide range of intensity of reproductive seasonality (r_birth_=[0.10-0.41]), with significant (Moremi) or non-significant (Tsaobis) birth peaks, and to some extent, various timings in their birth peaks (Moremi: around November, versus Drakensberg: around September). However, such population-level flexibility is often difficult to assess in many species, given the datasets available (*Papio ursinus* is indeed the only species in our sample represented by more than three populations with a reasonable number of births).

Flexibility in reproductive phenology may be facilitated by several mechanisms. First, a slower life history may allow papionin species to spread the energetic costs of reproduction over a prolonged period, such that pregnant or lactating females face only a small daily extra energetic expenditure that can be afforded at any time, as suggested by a recent modelling study (Burtschell et al., 2023). In addition, unlike many other mammals, cercopithecids do not use strict photoperiodic cues to trigger their reproduction (Heldstab et al., 2020) but may instead exhibit condition-induced reproduction, whereby conceptions (and/or cycle resumption after lactation) are more likely to occur when females are in better condition (Alberts et al., 2005; Beehner, Onderdonk, Alberts, & Altmann, 2006). Such reproductive flexibility may have contributed to their historical ecological success via their ability to colonize diverse environments, and may become a critical asset to facilitate their resilience to climate change, associated with increasing environmental unpredictability (Feng et al., 2013).

### Environmental unpredictability may drive flexible reproductive seasonality

We examined several climatic correlates of the intensity of reproductive seasonality across our sample, and found that environmental unpredictability was a significant predictor, with higher between-year rainfall variation being associated with lower reproductive seasonality. So far, most studies investigating climatic effects on reproductive phenology have focused on environmental seasonality, i.e., the magnitude of within-year environmental variation. In primates, the effect of environmental unpredictability on reproductive seasonality, e.g., through climatic events such as el Niño or fruit mast years in South-East Asia, has been suggested (Brockman & van Schaik, 2005a; van Schaik & van Noordwijk, 1985; Wiederholt & Post, 2011) but had never been tested. In line with our results, a previous study across 70 ungulate populations showed that the birth peak is more spread out in environments with more year-to-year environmental variation (English et al., 2012).

Unpredictable climates may considerably reduce the fitness benefits associated with seasonal breeding, such as enhancing maternal condition and offspring survival. In Africa, year-to-year climatic variation frequently takes the shape of an absence of rain during the rainy season (Alberts et al., 2005), which could cause severe reproductive costs in seasonal breeders who often synchronize lactation or weaning with the rainy season, subsequently forcing females to wait until the next breeding season to initiate a new reproductive event. In such conditions, other adaptive traits may be more advantageous than seasonal breeding to face the energetic costs of reproduction, such as the capacity to store energy (Brockman & van Schaik, 2005a; van Schaik & van Noordwijk, 1985), to expand the dietary repertoire via a generalist diet or foraging innovations (Grueter, 2017), to increase daily foraging time (Alberts et al., 2005; Grueter, 2017), to flexibly adjust lactation duration (Dezeure et al., 2021; van Noordwijk, 2012), or to reproduce cooperatively (Cornwallis et al., 2017; Lukas & Clutton-Brock, 2017). Papionin species show many such traits: they can store energy, they have an eclectic omnivorous diet, and they are flexible foragers that typically rely on fallback foods during the dry season (J. Altmann, Schoeller, Altmann, Muruthi, & Sapolsky, 1993; S. A. Altmann, 2009; Swedell, 2011). As such, it is likely that rainfall unpredictability selected these traits in papionin species (energy storage, omnivorous diet, slow life histories, etc.), which in turn contributed to shape their flexible reproductive seasonality.

Two recent studies found no or little effect of climatic unpredictability on the intensity of reproductive seasonality at the population level, one in *Papio ursinus* (Dezeure et al., 2023) and one on *Papio cynocephalus* using a modelling approach (Burtschell et al., 2023), questioning the robustness of the effect found in this study. This discrepancy may come from the fact that unpredictable climates select for reproductive flexibility, rather than nonseasonal breeding. In fact, the study by Burtschell et al., 2023 revealed that increasing climatic unpredictability was associated with a lower variance, but not a lower mean, in the intensity of reproductive seasonality. In addition, the effect of climatic unpredictability may be better detected across space than time, i.e., by comparing different populations living in distinct climates and environments, as is the case here, than by comparing the same population across time, as was the case for Dezeure et al., 2023. Different study designs should be combined, across time, taxonomic or spatial scales, to reveal the full complexity of selective pressures at play.

The numerous pressures affecting the intensity of reproductive seasonality are often hard to disentangle, meaning that the effects uncovered by correlational studies like ours may sometimes reflect other co-varying pressures, and should thus be interpreted cautiously. Specifically, environmental productivity, which is negatively correlated with climatic unpredictability at our study sites (cor=-0.80, t=-5.48, p<10^-4^), was also a significant predictor of the intensity of reproductive seasonality, but in a direction opposed to our prediction, as well as to previous results (Burtschell et al., 2023). In our sample, the least productive climates are also the most unpredictable, and environmental productivity may thus represent a confounding factor in the relationship between climatic unpredictability and seasonal breeding. In species with a flexible phenology, females can start a new reproductive event rapidly after a reproductive failure, without having to wait the next mating season. Such failures are likely to be particularly frequent where environmental productivity is low, contributing to spread reproductive events across the year cycle, and explaining how low environmental productivity may contribute to decrease reproductive seasonality. In addition, this study is based on datasets with heterogenous resolutions, including diverse numbers of births and years of study (e.g. Amboseli: N=496, Nyears=33, versus Queen Elizabeth: N=35, Nyears=2). Additional birth records in small datasets may change r_birth_, and could thus alter some of the results in our study. Lastly, our decomposition of rainfall components further calls for a more rigorous definition of the term seasonality, especially when it is used in a quantitative way. Indeed, a ‘more seasonal’ environment can either be an environment with higher within-year variation (i.e. the amplitude of variation between the ‘best’ and the ‘worst’ month of the year), with higher within-year over between-year variation (i.e. the amplitude of within-year variation controlling for the intensity of unpredictable variations), with a shorter productive season (i.e. the rainfall peak breadth), or with only a unimodal season (i.e. one rainy, or one warm season per year). Similarly, climatic unpredictability can be broken into two components: (1) the amount/magnitude of year-to-year variation (i.e., if the rainy season brings more or less rainfall than usual), and (2) year-to-year variation in the timing of the rainy season (i.e., if the rainy season occurs earlier or later than usual) (Clauss, Zerbe, Bingaman Lackey, Codron, & Müller, 2020). These various components have rarely been disentangled in empirical studies so far, and this study opens new methodological avenues to investigate various environmental components that are likely to affect reproductive seasonality.

### Females can match different reproductive stages with the food peak

The main pattern emerging from our investigations suggests that females from species with high breeding seasonality match lactation with the peak in vegetation productivity (Table 3). For species and populations with a more flexible reproductive phenology, females can match different reproductive stages with the annual vegetation peak, with a possible preference towards conception. This trend may reflect the condition-dependence of conception – a proximate mechanism – rather than an adaptive, optimal strategy aimed at synchronizing the vegetation peak with a particular reproductive stage. These results, obtained from (mostly) tropical primates, echo the broader mammalian literature showing that most mammals from temperate regions match lactation with the best season of the year (Bronson, 2009; Bronson & Heideman, 1994), while patterns are more variable in tropical and long-lived mammals, depending on body size, energy storage capacities and environmental predictability (Brockman & van Schaik, 2005a; Janson & Verdolin, 2005; van Schaik & van Noordwijk, 1985). In addition, even though weaning is a vulnerable life-history stage in young primates, which can be buffered when matched with the vegetation peak in a wild *Papio ursinus* population (Dezeure et al., 2021), few populations seemed to adopt this strategy.

Several caveats apply to the test of H2. First, although NDVI is a relatively good measure of plant productivity, highest values of NDVI do not necessarily coincide with the annual food peak, especially when focusing on omnivorous/frugivorous species. Precise phenological data from each population would be more accurate to quantify the annual food peak. Second, our estimations of lactation peak and weaning might lack accuracy, due to strong between-populations and between-individuals variation. Data quantifying maternal energy expenditure during lactation (Rosetta, Lee, & Garcia, 2011), or isotopic measures of trophic levels between mothers and infants (Carboni, Dezeure, Cowlishaw, Huchard, & Marshall, 2022; Reitsema, 2012) would be necessary, for each population, to determine the dynamics of lactation and weaning. Finally, additional unexplored factors can potentially affect reproductive timing and further limit our ability to detect a clear pattern. For example, for populations living at high altitudes like *Theropithecus gelada* from Simien and the *Papio ursinus* from Drakensberg, seasonal variation in temperatures also constrain reproductive phenology (Lycett et al., 1999; Tinsley Johnson et al., 2018).

### The evolution of reproductive flexibility in Anthropoid primates may inform our understanding of the reproductive phenology in early humans

Baboons and relatives represent an interesting model for understanding the evolution of behavioural and reproductive plasticity of early humans (J. Fischer et al., 2019; King, 2022). Although most great apes are nonseasonal breeders (Brockman & van Schaik, 2005b; Campos et al., 2017), suggesting that their common ancestor also bred year-round, humans are distinct from other apes by exhibiting much faster reproductive paces (which are similar to most Africa-dwelling papionins: Swedell, 2011), and by living in a wider variety of environments, rather than being restricted to tropical forests (Wells & Stock, 2007). The selective pressures that have shaped reproductive phenology in the human lineage versus in other apes may therefore differ, and the papionins, who have similarly left forested habitats to colonize savannahs may provide valuable insights to understand the adaptation of early humans to such diverse and unpredictable environments. Our results suggest that baboons have acquired a low and flexible reproductive seasonality, as well as several other adaptive traits, when facing more arid and unpredictable environments, such as a generalist omnivorous diet and the frequent use of fallback foods, frequent foraging innovations, an increased ability to store fat and to switch home ranges to more suitable areas. Similar reproductive, physiological and behavioural adaptations to environmental unpredictability may have allowed early humans to thrive and maintain fast reproductive paces during their colonization of a wide variety of environments.

## Conclusion

Our work revealed substantial variation in patterns of reproductive seasonality within and across species of Africa-dwelling papionins, highlighting an exceptional flexibility in their reproductive phenology. Among multiple dimensions of climatic variation, rainfall unpredictability and productivity were the main predictors of the intensity of reproductive seasonality, with arid and unpredictable climates being associated with less seasonal reproduction. Among populations with a pronounced breeding seasonality, females often match lactation with the annual vegetation peak, while phenology patterns are very diverse in other populations. This study sheds new light on the selective pressures shaping reproductive seasonality in long-lived tropical mammals, as well as on potential adaptations to environmental unpredictability. It may further provide an original contribution to understand why humans breed year-round, given their phylogenetic ties and convergences in life-history and ecology with our taxonomic group.

## ACKNOWLEDGMENTS

We thank Cassandra Raby for her kind help on extracting NDVI data, and Christophe Dezeure for making the Figure 1 of this paper (and more generally for his paternal care). This project was funded by a grant from the Agence Nationale de la Recherche (ANR ERS-17-CE02-0008, 2018-2021) awarded to E. H.

## DATA AVAILABILITY

The raw data and scripts used for this paper are available in the following Zenodo repository: <u>10.5281/zenodo.13312416</u>.

