## Supplementary Material for "Flexible reproductive seasonality in Africa-dwelling papionins is associated with low environmental productivity and high climatic unpredictability"

### **Appendix S1. Comparisons of measures of the intensity of birth seasonality.**

In order to select one or more measure of the intensity of birth seasonality, we computed four classical measures (see Table S1):

(i) *The length of the r-vector* *(r_birth_)* measures the degree of uniformity of the birth distribution across the annual cycle and varies from 0 to 1 (Di Bitetti & Janson, 2000; Janson & Verdolin, 2005; Thompson & McCabe, 2013). When r_birth_=0, births are evenly spread across months (i.e. non-seasonal), while when r_birth_=1, births all occur at the exact same month of the year (extremely seasonal).
(ii) *The breadth of the birth peak* *in a given population using the minimum number of consecutive months in which 50% of births occurred* (called ‘BPB 50’).
(iii) *The breadth of the birth peak using the minimum number of consecutive months in which 80% of births occurred* (called ‘BPB 80’) (Heldstab et al., 2018, 2020; Zerbe et al., 2012).
(iv) *A categorical measure of reproductive seasonality, i.e. significant vs. non-significant birth peak.* To do so, we ran Rayleigh tests to assess, statistically, if birth distributions are uniform (H_0_: r_birth_ =0) or not (H_1_: r_birth_ ≠0) along the annual cycle, using the ‘r.test’ function from ‘CircStats’ package (Agostinelli & Lund, 2018). Populations’ birth peaks were categorized as significant when the result of the Rayleigh test allowed us to reject the null hypothesis of a uniform birth distribution.

We found strong correlations between r_birth_ and both BPB 50 (cor= -0.91, p<10^-4^) and BPB 80 (cor= -0.99, p<10^-4^), and thus opted to use r_birth_ in all downstream analyses.

### **Appendix S2. Method to compute the between-year variation in the timing of rainfall**

We were interested to test the effect of the unpredictability of the rainfall peak in term of timing on the intensity of reproductive seasonality (H1-7), and we thus had to quantify how much the timing of rainfall varied between-years. First, for each population, we used the predictable rainfall variation component (K_rain_+Rainfall S) to compute the mean rainfall date µ_rain_, which indicates the annual peak of rainfall. To do so, we used circular statistics: the length of the vector for each month equals the amount of rainfall (in mm) that fell this month, and then computed (similarly to what we did for µ_birth_) µ_rain_, as the angle, converted in a date, of the r-vector of rainfall values (Markham, 1970). We used the function ‘circ.summary’ from the ‘CircStats’ package (Agostinelli & Lund, 2018), and the same methodology allowed us to compute µ_NDVI_. Second, we computed the mean rainfall date for each year. We separated our rainfall data in 22 distinct periods of 365 days centered on µ_rain_. For example, if µ_rain_ =10^th^ of September, periods ran from March, 10th to the next March, 10th. We similarly separated our rainfall data in 44 distinct semesters (n=365/2 days) for environments with 2 rainy seasons: the first semester going from February to July, and the second one going from August to January, given the observed distribution of the seasonal component of rainfall in these environments (see Figure 2). Third, for each given year (or semester), we computed the mean rainfall date using circular statistics. Fourth, we computed the standard deviation of these mean dates per year and population, to quantify the between-year rainfall variation in timing, with higher values corresponding to higher uncertainty in the annual timing of rainfall. For environments with 2 rainy seasons, we quantified two standard deviations, one per semester, and further computed the mean of these two standard deviations as the measure of between-semester rainfall variation in timing.

### **Appendix S3. Definition of the targeted reproductive windows used in tests of Hypothesis 2, regarding the reproductive stage synchronized with the food peak**

For H2-1 (‘conception’), we first determined the mean conception date per population with the formula: µ_conc_ = µ_birth_ – gestation length. We used the mean gestation length (in days) per population when it was provided in the literature, or used a species-specific gestation length otherwise, because within-species variation in gestation length is low (see Table S3). We tested if females tended to conceive during, soon before, or soon after the annual food peak, comparing random NDVI values to observed seasonal NDVI values in (i) the six months surrounding µ_conc_, (ii) the three months before µ_conc_ and (iii) the three months after µ_conc_. For H2-2 (‘lactation’), we considered lactation period as the 6 months following birth. Indeed, this window has often been used to characterize lactation and infant dependency, and in particular the period of high infanticide risk, in several baboon populations (Altmann, 1980; Palombit, 2003). We tested if females tended to adjust the entire lactation period, early-lactation (3 months post-birth) or mid-lactation (3-6 months post-birth) with the annual food peak, by comparing random NDVI values to seasonal NDVI values in the 6 months after µ_birth_, the 3 months after µ_birth_ or from 3 to 6 months after µ_birth,_ respectively. For H2-3 (‘weaning’), we determined the mean weaning age per population using as a proxy the mean duration of post-partum amenorrhea (‘PPA’, in days) (Borries, Lu, Ossi-Lupo, Larney, & Koenig, 2014; Lee, Majluf, & Gordon, 1991) (see Table S3). We computed the mean weaning date, µ_PPA_, as µ_birth_ + PPA. We tested if females tended to adjust weaning, mid-weaning (here approximated by the 3 months before µ_PPA_) or late-weaning (here approximated by the 3 months following µ_PPA_) with the annual food peak, by comparing random NDVI values to seasonal NDVI values observed in the 6 months surrounding µ_PPA_, and in the 3 months before and after µ_PPA_, respectively.

### **Table S1:** Birth seasonality data for the populations considered in this study

| **Species** | **Population** | **Number of births** | **Number of years of survey** | **r_birth_** | **P-value  (Rayleigh)** | **BPB**  **80**  **(months)** | **BPB**  **50**  **(months)** | **µ_birth_** | **References** |
| --- | --- | --- | --- | --- | --- | --- | --- | --- | --- |
| *Cercocebus  atys* | Taï | 52 | 3 | 0.8312 | <10^-5^ | 3 | 2 | 5-Jan | (Range, Förderer, Storrer-Meystre, Benetton, & Fruteau, 2012) |
| *Cercocebus  sanjei* | Udzungwu  Mountains | 28 | 3 | 0.5588 | <10^-5^ | 6 | 3 | 17-Aug | (Thompson & McCabe, 2013) |
| *Lophocebus  albigena* | Kibale | 72 | 9 | 0.2762 | 0.0041 | 8 | 5 | 13-Feb | (Arlet et al., 2014) |
| *Macaca  sylvanus* | Akfada | 56 | 8 | 0.9350 | <10^-5^ | 2 | 2 | 6-May | (Ménard & Vallet, 1993) |
|  | Tigounatine | 75 | 8 | 0.9467 | <10^-5^ | 2 | 1 | 24-May |  |
| *Mandrillus  sphinx* | Lekedi | 218 | 9 | 0.6766 | <10^-5^ | 5 | 2 | 1-Jan | (Dezeure, Charpentier, & Huchard, 2022) |
|  | Moukalaba- Doudou | 208 | 2 | 0.7940 | <10^-5^ | 3 | 2 | 8-Jan | (Hongo, Nakashima, Akomo-Okoue, & Mindonga-Nguelet, 2016) |
| *Papio  anubis* | Gashaka-  Gumti | 32 | 5 | 0.2167 | 0.2237 | 8 | 4 | 17-Dec | (Higham et al., 2009) |
|  | Gilgil | 118 | 10 | 0.1873 | 0.0159 | 9 | 5 | 23-Feb | (Bercovitch & Harding, 1993) |
|  | Queen  Elizabeth | 35 | 2 | 0.0971 | 0.7216 | 9 | 5 | 17-Jan | (Rowell, 1966) |
| *Papio  cynocephalus* | Amboseli | 496 | 33 | 0.1344 | 0.0001 | 9 | 6 | 12-Oct | (Alberts et al., 2005) |
|  | Mikumi | 164 | 8 | 0.1584 | 0.0163 | 9^+^ | 6^+^ | 24-Jul | (Rhine, Wasser, & Norton, 1988) |
|  | Tana  River | 35 | 5 | 0.1413 | 0.5004 | 9 | 6 | 25-Nov | (Bentley-Condit & Smith, 1997) |
| *Papio  hamadryas* | Filoha | 218 | 7 | 0.0208 | 0.9097 | 10 | 6 | 21-Jun | Swedell, unpubl. data |
| *Papio  kindae* | Kasanka | 70 | 7 | 0.4971 | <10^-5^ | 6 | 4 | 16-Jul | (Petersdorf, Weyher, Kamilar, Dubuc, & Higham, 2019) |
| *Papio  ursinus* | De Hoop | 30 | 4 | 0.0969 | 0.7574 | 9 | 5 | 20-Oct | (Barrett, Henzi, & Lycett, 2006) |
|  | Drakensberg | 37 | 7 | 0.4143 | 0.0014 | 6 | 4 | 23-Nov | (Lycett, Weingrill, & Henzi, 1999) |
|  | Moremi | 122 | 10 | 0.3710 | <10^-5^ | 7 | 4 | 27-Sep | (Cheney et al., 2004) |
|  | Tokai | 52 | 2 | 0.2394 | 0.0507 | 8 | 5 | 21-Dec | Chowdhury, unpubl. data |
|  | Tsaobis | 215 | 15 | 0.1046 | 0.0949 | 9 | 5 | 18-Nov | (Dezeure et al., 2021) |
| *Theropithecus  gelada* | Simien | 354 | 9 | 0.1550 | 0.0002 | 9 | 5 | 8-Oct | (Tinsley Johnson, Snyder-Mackler, Lu, Bergman, & Beehner, 2018) |

^+^ Approximated birth peak breadths for Mikumi, given that the number of births was only provided per 3 month-period (not per month)

Populations with significant birth peak (P-values of the Rayleigh test <0.05) are grey shaded.

### **Table S2:** Components of environmental variation per population

These data have been extracted and computed following the methods described in the main text (see 3-Environmental data).

| **Species** | **Population** | **Country** | **GPS coordinates  (West,South, East,North)** | **Latitude (°)** | **Mean  annual  rainfall (mm)** | **Magnitude of  rainfall seasonality** | **Number of  rainy  season(s)** | **Rainfall peak breadth (months)** | **Magnitude of rainfall unpredictability** | **Timing of rainfall unpredictability (days)** | **Habitat** | **µ_rain_** | **µ_NDVI_** |
| --- | --- | --- | --- | --- | --- | --- | --- | --- | --- | --- | --- | --- | --- |
| *Cercocebus  atys* | Taï | Ivory  Coast | 7.26, 5.75,  -7.21, 5.80 | 5.8 | 1691 | 3.73 | 2 | NA | 3.08 | 9.31 | Tropical  forest | 18-Jul | 6-Feb |
| *Cercocebus  sanjei* | Udzungwu  Mountains | Tanzania | 36.80, -7.78,  36.88, -7.68 | 7.7 | 1187 | 7.45 | 1 | 5 | 3.98 | 14.96 | Tropical  forest | 20-Feb | 30-Sep |
| *Lophocebus  albigena* | Kibale | Uganda | 30.40, 0.45, 30.45, 0.50 | 0.5 | 1155 | 3.76 | 2 | NA | 2.99 | 8.49 | Tropical  forest | 30-Sep | 19-May |
| *Macaca  sylvanus* | Akfadou | Algeria | 36.69, 4.55, 36.71, 4.61 | 36.7 | 760 | 4.03 | 1 | 8 | 5.82 | 33.85 | Mosaic  forest-grassland | 12-Jan | 2-Aug |
|  | Tigounatine | Algeria | 36.4, 4.13, 36.45, 4.18 | 36.4 | 789 | 4.11 | 1 | 8 | 5.71 | 33.81 | Mosaic  forest-grassland | 18-Jan | 15-Mar |
| *Mandrillus  sphinx* | Lekedi | Gabon | 13.03, -1.81,  13.04, -1.80 | 1.8 | 1913 | 4.96 | 1 | 7 | 3.17 | 12.16 | Tropical forest | 12-Jan | 11-Apr |
|  | Moukalaba- Doudou | Gabon | 10.24,-2.60, 10.35,-2.50 | 2.5 | 1482 | 6.26 | 1 | 6 | 3.76 | 13.37 | Tropical forest | 20-Jan | 30-Mar |
| *Papio  anubis* | Gashaka- Gumti | Nigeria | 11.58,7.51, 11.66,7.56 | 7.5 | 1969 | 7.06 | 1 | 6 | 3.50 | 8.88 | Mosaic  forest-grassland | 31-Jul | 5-Aug |
|  | Gilgil | Kenya | 36.23,-0.61,  36.45, -0.48 | 0.5 | 1220 | 2.62 | 2 | NA | 3.79 | 10.29 | Open  savannah | 29-Apr | 23-Jun |
|  | Queen  Elizabeth | Uganda | 29.70, -0.57,  29.80, -0.47 | 0.5 | 1015 | 2.99 | 2 | NA | 3.21 | 9.00 | Mosaic  forest-grassland | 6-Nov | 8-Dec |
| *Papio  cynocephalus* | Amboseli | Kenya | 37.27, -2.71,  37.32, -2.62 | 2.7 | 490 | 7.77 | 2 | NA | 8.93 | 10.29 | Open  savannah | 31-Jan | 14-Mar |
|  | Mikumi | Tanzania | 37.37, -7.29,  37.49, -7.19 | 7.2 | 1296 | 4.43 | 1 | 8 | 4.87 | 46.98 | Open  savannah | 25-Feb | 13-Apr |
|  | Tana River | Kenya | 40.13, -1.94,  40.13, -1.93 | 1.9 | 608 | 6.04 | 2 | NA | 7.64 | 10.78 | Open  savannah | 12-Jan | 4-Mar |
| *Papio  hamadryas* | Awash | Ethiopia | 39.96, 9.00,  40.07, 9.12 | 9.05 | 762 | 7.36 | 1 | 6 | 4.92 | 12.95 | Open  savannah | 22-Jul | 23-Aug |
| *Papio  kindae* | Kasanka | Zambia | 30.21,-12.58, 30.22,-12.55 | 12.6 | 1148 | 9.73 | 1 | 5 | 4.16 | 7.11 | Mosaic  forest-grassland | 19-Jan | 26-Feb |
| *Papio  ursinus* | De Hoop | South  Africa | 20.52,-34.45, 20.60,-34.39 | 34.4 | 373 | 2.08 | 1 | 9 | 8.82 | 94.32 | Open  savannah | 17-Jun | 29-May |
|  | Drakensberg | South  Africa | 29.44,-29.28,  29.54, -29.15 | 29.2 | 946 | 6.30 | 1 | 6 | 3.89 | 13.22 | Open  savannah | 12-Jan | 5-Feb |
|  | Moremi | Botswana | 22.86, -19.42,  23.01, -19.27 | 19.3 | 507 | 7.63 | 1 | 5 | 7.25 | 22.91 | Open  savannah | 26-Jan | 20-Feb |
|  | Tokai | South Africa | 18.39, -34.08,  18.43, -34.05 | 34.1 | 517 | 4.46 | 1 | 8 | 7.41 | 22.79 | Mosaic  forest-grassland | 5-Jul | 22-Jun |
|  | Tsaobis | Namibia | 15.67, -22.46,  15.88, -22.30 | 22.4 | 239 | 9.87 | 1 | 4 | 13.65 | 18.29 | open  savannah | 19-Feb | 9-Apr |
| *Theropithecus  gelada* | Simien | Ethiopia | 38.31, 13.20,  38.42, 13.28 | 13.2 | 1007 | 6.63 | 1 | 7 | 4.28 | 16.03 | Open  savannah | 24-Jul | 23-Aug |

### **Table S3**: Reproductive parameters per population.

| Species | Population | Gestation   (days) | Weaning age (days) | References |
| --- | --- | --- | --- | --- |
| *Cercocebus  atys* | Taï (reproductive data from a  captive population) | 167 | 216 | (Gust, Busse, & Gordon, 1990) (captive population) |
| *Cercocebus  sanjei* | Udzungwu  Mountains | 172 | 204 | (Fernández, Doran-Sheehy, Borries, & Brown, 2014) |
| *Lophocebus  albigena* | Kibale | 186 | 224 | (Wallis, 1983) |
| *Macaca sylvanus* | Akfadou | 164* | NA | (Kingdon et al., 2012) |
|  | Tigounatine | 164* | NA | (Kingdon et al., 2012) |
| *Mandrillus  sphinx* | Lekedi | 175 | 288 | (Dezeure et al., 2022) |
|  | Moukalaba- Doudou | 175* | NA |  |
| *Papio  anubis* | Gashaka- Gumti | 185 | 322 | (Higham et al., 2009) |
|  | Gilgil | 180 | 407 | (Smuts & Nicolson, 1989) |
|  | Queen  Elizabeth | 180* | NA |  |
| *Papio  cynocephalus* | Amboseli | 178 | 332 | (Gesquiere, Altmann, Archie, & Alberts, 2017) |
|  | Mikumi | 178* | NA |  |
|  | Tana River | 182 | 443 | (Bentley-Condit & Smith, 1997) |
| *Papio  hamadryas* | Filoha | 182 | 266 | (Swedell, 2006, 2011) |
| *Papio  kindae* | Kasanka | 178* | NA |  |
| *Papio  ursinus* | De Hoop | 190 | 265 | (Weingrill, Gray, Barrett, & Henzi, 2004) |
|  | Drakensberg | 190* | NA |  |
|  | Moremi | 183 | 453 | (Cheney et al., 2004; Johnson & Bock, 2004) |
|  | Tokai | 190* | NA |  |
|  | Tsaobis | 190 | 353 | (Dezeure et al., 2021) |
| *Theropithecus  gelada* | Simien | 183 | 534 | (Roberts, Lu, Bergman, & Beehner, 2017) |

* Mean gestation length was not available for these populations. So, we used mean gestation length of other population of the same species with the highest sample size. We considered that the gestation length of Kinda baboons was similar to the ones of yellow baboons.

### **Table S4:** Reproductive timing in relation to food availability seasonality

For each population with a significant birth peak, we give the statistics and P-values of the Fisher-Pitman permutation test (Z test) investigating if NDVI (our proxy of food availability) is higher than random during different stages of female reproductive cycle, namely conception, lactation, or weaning. For the conception hypothesis (H2.1), we looked at the NDVI values either around (3 months before and after), 3 months before or 3 months after the mean conception date µ_conc_. For the lactation hypothesis (H2.2), we looked at NDVI values during either the 6 months following the mean birth date µ_birth_ (‘Whole lactation’), the 3 months after µ_birth_ (‘Early lactation’), or from 3 to 6 months after µ_birth_ (‘Mid lactation’). For the weaning hypothesis (H2.3), we looked at the NDVI values either around (3 months before and after µ_PPA_), 3 months before or 3 months after the mean weaning date µ_PPA_. Significant effects are indicated in bold. In addition, we indicated the difference, in months, between the mean conception date (µ_conc_) and the mean NDVI date (µ_NDVI_), between the mean NDVI date (µ_NDVI_) and the mean birth date (µ_birth_), and between the mean NDVI date (µ_NDVI_) and the mean weaning date (µ_PPA_).

| Species | Population | ***Conception hypothesis (H2.1)*** | | | | | | |
| --- | --- | --- | --- | --- | --- | --- | --- | --- |
|  |  | Around conception | | Before conception | | After conception | | µ_conc_- µNDVI (month) |
|  |  | Z test | P-value | Z test | P-value | Z test | P-value |  |
| *Cercocebus  atys* | Taï | 2.642 | 1.000 | 1.492 | 0.932 | 1.559 | 0.941 | 5.45 |
| *Cercocebus  sanjei* | Udzungwa | 0.517 | 0.694 | **-1.573** | **0.045** | 2.170 | 0.986 | 4.91 |
| *Lophocebus  albigena* | Kibale | -1.00 | 0.167 | **-2.262** | **0.009** | 1.107 | 0.855 | 2.73 |
| *Macaca  sylvanus* | Akfadou | 1.187 | 0.876 | -1,082 | 0.182 | 2.452 | 0.991 | 3.71 |
|  | Tigounatine | 0.780 | 0.771 | 1.096 | 0.859 | -0.755 | 0.240 | -3.09 |
| *Mandrillus  sphinx* | Lekedi | -0.616 | 0.277 | **-1.772** | **0.032** | 1.061 | 0.841 | 2.93 |
|  | Moukalaba | 1.585 | 0.947 | -1.081 | 0.177 | 2.911 | 1.000 | 3.55 |
| *Papio  anubis* | Gilgil | -0.514 | 0.314 | -0.994 | 0.186 | 0.400 | 0.627 | 2.10 |
| *Papio  cynocephalus* | Amboseli | -1.616 | 0.056 | **-1.701** | **0.05** | -0.164 | 0.459 | 1.12 |
|  | Mikumi | -0.744 | 0.242 | 0.599 | 0.732 | -1.458 | 0.077 | -2.5 |
| *Papio  kinda* | Kasanka | **-2.707** | **0.002** | -1.03 | 0.168 | **-2.096** | **0.005** | -1.28 |
| *Papio  ursinus* | Drakensberg | 1.209 | 0.881 | -0.897 | 0.177 | 2.294 | 0.996 | 3.35 |
|  | Moremi | **-2.312** | **0.012** | **-2.649** | **0.005** | -0.021 | 0.468 | 1.25 |
| *Theropitethecus gelada* | Simien | 2.291 | 0.994 | 2.201 | 1.000 | 0.445 | 0.677 | -4.50 |
|  |  | ***Lactation hypothesis (H2.2)*** | | | | | | |
|  |  | Whole lactation | | Early lactation | | Mid lactation | | µ_NDVI_- µ_birth  (month)_ |
|  |  | Z test | P-value | Z test | P-value | Z test | P-value |  |
| *Cercocebus  atys* | Taï | **-1.607** | **0.043** | **-1.775** | **0.009** | -0.081 | 0.482 | 1.05 |
| *Cercocebus  sanjei* | Udzungwa | -0.685 | 0.254 | 0.782 | 0.773 | **-1.573** | **0.045** | 1.45 |
| *Lophocebus  albigena* | Kibale | -1.474 | 0.070 | 0.561 | 0.713 | **-2.262** | **0.009** | 3.16 |
| *Macaca  sylvanus* | Akfadou | **-2,604** | **0.001** | **-1.773** | **0.009** | -1.124 | 0.136 | 2.89 |
|  | Tigounatine | 2.413 | 0.993 | 1.690 | 0.955 | 1.096 | 0.859 | -2.30 |
| *Mandrillus  sphinx* | Lekedi | **-2.432** | **0.005** | -1.036 | 0.173 | **-1.772** | **0.032** | 3.32 |
|  | Moukalaba | **-2.405** | **0.006** | -1.474 | 0.064 | -1.304 | 0.114 | 2.70 |
| *Papio  anubis* | Gilgil | -1.382 | 0.089 | -0.602 | 0.286 | -0.994 | 0.186 | 3.98 |
| *Papio  cynocephalus* | Amboseli | -0.439 | 0.334 | 0.743 | 0.768 | -1.25 | 0.109 | 5.03 |
|  | Mikumi | 2.635 | 0.999 | 2.443 | 0.996 | 0.599 | 0.732 | 8.65 |
| *Papio  kinda* | Kasanka | 1.419 | 0.920 | 2.669 | 1.000 | -1.03 | 0.168 | 7.36 |
| *Papio  ursinus* | Drakensberg | **-2.694** | **0.002** | **-2.213** | **0.009** | -0.897 | 0.177 | 2.40 |
|  | Moremi | **-1.838** | **0.031** | 0.528 | 0.691 | **-2.649** | **0.005** | 4.77 |
| *Theropitethecus gelada* | Simien | 1.922 | 0.974 | 0.019 | 0.482 | 2.201 | 1.000 | -1.51 |
|  |  | ***Weaning hypothesis (H2.3)*** | | | | | | |
|  |  | Around weaning | | Before weaning | | After weaning | | µ_NDVI_- µ_PPA  (month)_ |
|  |  | Z test | P-value | Z test | P-value | Z test | P-value |  |
| *Cercocebus  atys* | Taï | 2.642 | 1.000 | 1.492 | 0.932 | 1.559 | 0.941 | 5.95 |
| *Cercocebus  sanjei* | Udzungwa | 0.517 | 0.694 | **-1.573** | **0.045** | 2.170 | 0.986 | -5.26 |
| *Lophocebus  albigena* | Kibale | 0.807 | 0.775 | -0.64 | 0.296 | 1.571 | 0.932 | -4.21 |
| *Macaca  sylvanus* | Akfadou | NA | NA | NA | NA | NA | NA | NA |
|  | Tigounatine | NA | NA | NA | NA | NA | NA | NA |
| *Mandrillus  sphinx* | Lekedi | 2.797 | 1.000 | 2.324 | 0.986 | 0.905 | 0.818 | 5.85 |
|  | Moukalaba | NA | NA | NA | NA | NA | NA | NA |
| *Papio  anubis* | Gilgil | -0.077 | 0.462 | 2.323 | 0.996 | **-2.411** | **0.005** | 2.62 |
| *Papio  cynocephalus* | Amboseli | 2.978 | 1.000 | 1.276 | 0.900 | 2.163 | 0.986 | -5.88 |
|  | Mikumi | NA | NA | NA | NA | NA | NA | NA |
| *Papio  kinda* | Kasanka | NA | NA | NA | NA | NA | NA | NA |
| *Papio  ursinus* | Drakensberg | NA | NA | NA | NA | NA | NA | NA |
|  | Moremi | **-1.838** | **0.031** | 0.528 | 0.691 | **-2.649** | **0.005** | 1.87 |
| *Theropitethecus gelada* | Simien | 2.291 | 0.994 | 2.201 | 1.000 | 0.445 | 0.677 | 4.95 |

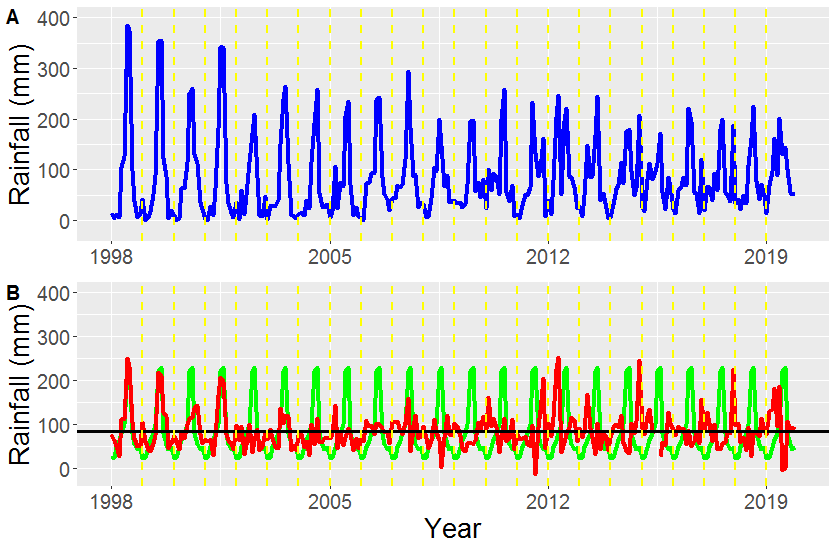

### **Figure S1**: Example of rainfall variation decomposition

We represented in Panel A the average monthly cumulative rainfall (raw data, in mm) recorded at Simien National Park (example from the gelada population) over 22 years (from January 1998 to December 2019) in blue. In Panel B, the black horizontal line indicates the mean monthly rainfall (K_rain_), the green curve represents the predictable (seasonal, i.e. repeatable pattern between years) rainfall variation (K_rain_ + Rainfall S), and the red curve represents the unpredictable (between-year) rainfall variation (K_rain_ + Rainfall NS) over 22 years of records.
